# Engraftment of donor phageome via fecal microbiota transplantation in recurrent *C. difficile* infection: a prospective observational study

**DOI:** 10.1101/2025.10.09.681403

**Authors:** Carmen Chen, Trestan Pillonel, Alessia Carrara, Julien Schaer, Gregory Resch, Tatiana Galperine, Benoit Guery, Claire Bertelli

**Author notes:** These authors contributed equally to this work (Benoit Guery, Claire Bertelli).

## Abstract

The high efficacy of fecal microbiota transplantation (FMT) in treating recurrent *Clostridioides difficile* infection (rCDI) is often attributed to the restoration of the bacterial community. However, factors beyond bacteria, such as bacteriophages (phages), may also play a critical role in FMT’s success.

We aimed to evaluate the preservation of the phage community (phageome) along the FMT production process following Good Manufacturing Practices (GMP) at the Lausanne University Hospital (CHUV), and the engraftment of the phages in patients receiving FMT to treat rCDI. Samples from one donor were used to test the need for amplification and to compare spin-column versus magnetic bead purification. Then, sixteen samples, from four donations of a second healthy donor, were collected at various production stages – fresh, frozen, homogenized, and encapsulated – for phageome analysis. The phage community profiles of three patients before, at 14, and 60 days after FMT were examined to evaluate donor phage engraftment.

Phages were detected in all sample types, and samples clustered by donation, indicating that the pre-processing steps did not significantly alter the phage profile. The recipients’ phageome prior to FMT was characterized by low diversity, each recipient being dominated by a different phage. In contrast, the profile 14 days post-FMT demonstrated the engraftment of donor-derived phages, which persisted at 60 days. Most were predicted to be temperate phages of the Caudoviricetes class infecting members of the Clostridia bacterial class, and *Lachnospiraceae* and *Oscillospiraceae* bacterial families.

Our findings suggest that the CHUV production process for oral FMT capsules preserves the phage community and that donor phages successfully engraft in recipients. Further larger-scale studies and intervention trials will help elucidate the mechanisms underlying the potential of phages in FMT’s efficacy.

## 2 Introduction

Infection by *C. difficile* can cause symptoms ranging from mild diarrhea to severe complications with toxic megacolon, colon perforation, septic shock, and organ failure^1^. Commonly found in the gastrointestinal tract of mammals and transmitted through the fecal-oral route, this bacterium is a major cause of nosocomial infections. Risk factors associated with *C. difficile* infection (CDI) include antibiotic exposure, older age, and hospitalization. Typically, high concentrations of secondary bile acids produced by the commensal bacteria of the gut microbiota prevent *C. difficile* spore germination. However, the loss of the bacterial community after antibiotic exposure promotes germination into vegetative cells and CDI^2^.

After a first episode of CDI, 10-25% of patients will experience a first recurrence. Among those who experienced a first recurrence, 65% will develop recurrent episodes^1^. At the Lausanne University Hospital (CHUV) in Switzerland, the standard of care for patients experiencing a second recurrence is fecal microbiota transplant (FMT) through frozen oral capsules. These capsules are produced at CHUV using stools obtained from healthy donors who have passed a stringent screening following international recommendations Appendix C. Although FMT treats CDI with a greater efficacy rate (80-90%)^3^ than antibiotics (20-40%)^4^, the precise mechanism leading to the success of FMT remains unclear. The most widely accepted explanation is that FMT restores the bacterial community lost due to antibiotic treatment^5^.

In a healthy condition, or eubiosis, the gut bacterial composition demonstrates high taxonomic diversity and a stable core microbiota. This healthy status can be disturbed through antibiotic treatment or pathogen invasion, such as CDI. Termed dysbiosis, the imbalance in the bacterial composition and the resulting alteration of metabolic activity has been shown to be associated with many diseases, including obesity, inflammatory bowel disease, Crohn’s disease, and celiac disease^6^.

Similar to gut bacterial imbalance, dysbiosis in the gut phageome, referring to the community of phages in the gut, has also been shown to be associated with disease^7^. There are approximately 10^9^ viral particles per gram of feces, of which more than 90% are phages^8,9^. These small viruses, which infect specific bacteria, have co-evolved with their hosts, shaping and promoting diversity within microbial communities. Links between viral and bacterial composition have been identified; bacterial abundance and diversity being positively correlated to the viral profile^10^. Thus, in addition to bacteria, a large number of phages are likely transferred during FMT, and these phages may be partially responsible for its effectiveness^8^.

Dysbiosis in the phageome is typically reflected by a loss of viral diversity^7^. The phage profile of CDI patients revealed a significantly higher abundance of *Caudovirales*, but lower *Caudovirales* diversity, richness, and evenness, compared to healthy individuals^11^. Patients with ulcerative colitis showed decreased *Caudovirales* diversity, richness, and evenness^12^, while patients with metabolic syndrome exhibited an enrichment of phages infecting *Streptococcaceae* and *Bacteroidaceae* and a depletion of phages infecting *Bifidobacteriaceae*^13^.

While the role of bacteria in FMT is widely accepted, the contribution of phages in FMT success has also been recently suggested. Fecal virome transplant (FVT) is the transplant of FMT devoid of all intact bacteria. Similarly to traditional FMT, FVT treatment in CDI was reported to restore normal stool habits and eliminated all symptoms, suggesting a potential role of phages or metabolites in mediating CDI, although these findings are based on a limited sample size^14^. Additionally, successful FMT treatments have shown engraftment of donor-derived phages, further supporting the idea that phages contribute to FMT efficacy^11^. To explore whether phages could also play a role in FMT success in rCDI patients at CHUV, this study investigated whether the pre-processing steps used to produce oral capsules altered the phage profile, assessed the engraftment of donor-derived phages in recipients, and evaluated their temporal stability.

## 3 Material & Methods

### 3.1 Sample collection

Two healthy volunteers (D01 and D02) who passed a stringent screening following international recommendations ^Appendix C^ to become fecal microbiota transplant (FMT) donors were recruited at CHUV (**Fig 1A**). D01 provided six donations between Jan-Feb 2022, while D02 contributed five donations between Oct-Nov 2023. To assess the preservation of the phage composition in CHUV’s FMT capsules, 16 samples from D02 were collected throughout the production process. Four recipients were recruited between May-Sep 2024; one was excluded due to missing samples. Nine samples from the remaining three recipients, collected before and after FMT, were used to evaluate donor phage engraftment and temporal stability (**Fig 1B**). These three patients had each experienced *≥* 2 rCDI episodes and subsequently received FMT. All patients had undergone multiple rounds of antibiotics for previous episodes and were pre-treated with vancomycin while awaiting FMT to prevent further recurrences, which was discontinued 48 hours before the procedure (**Table 1**). To establish baseline profiles, fresh samples before FMT (Fresh.d0) were collected. All three patients then received capsules produced from the same batch (D0205). The FMT regimen consisted of ingesting 2*×*20 FMT capsules within 48 hours. Follow-up samples were collected at 14 (Fresh.d14) and 60 (Fresh.d60) days post-FMT (dp-FMT) to investigate the impact of FMT on the viral community. All patients provided a signed informed consent, and this study was approved by the Ethics Committee of Vaud (CER-VD 2021-00674). Patients were considered cured if they remained symptom-free eight weeks post-FMT as recommended by ESCMID^15^. Further details on the recruitment process are provided in Appendix A and Appendix C.

**Figure 1:**
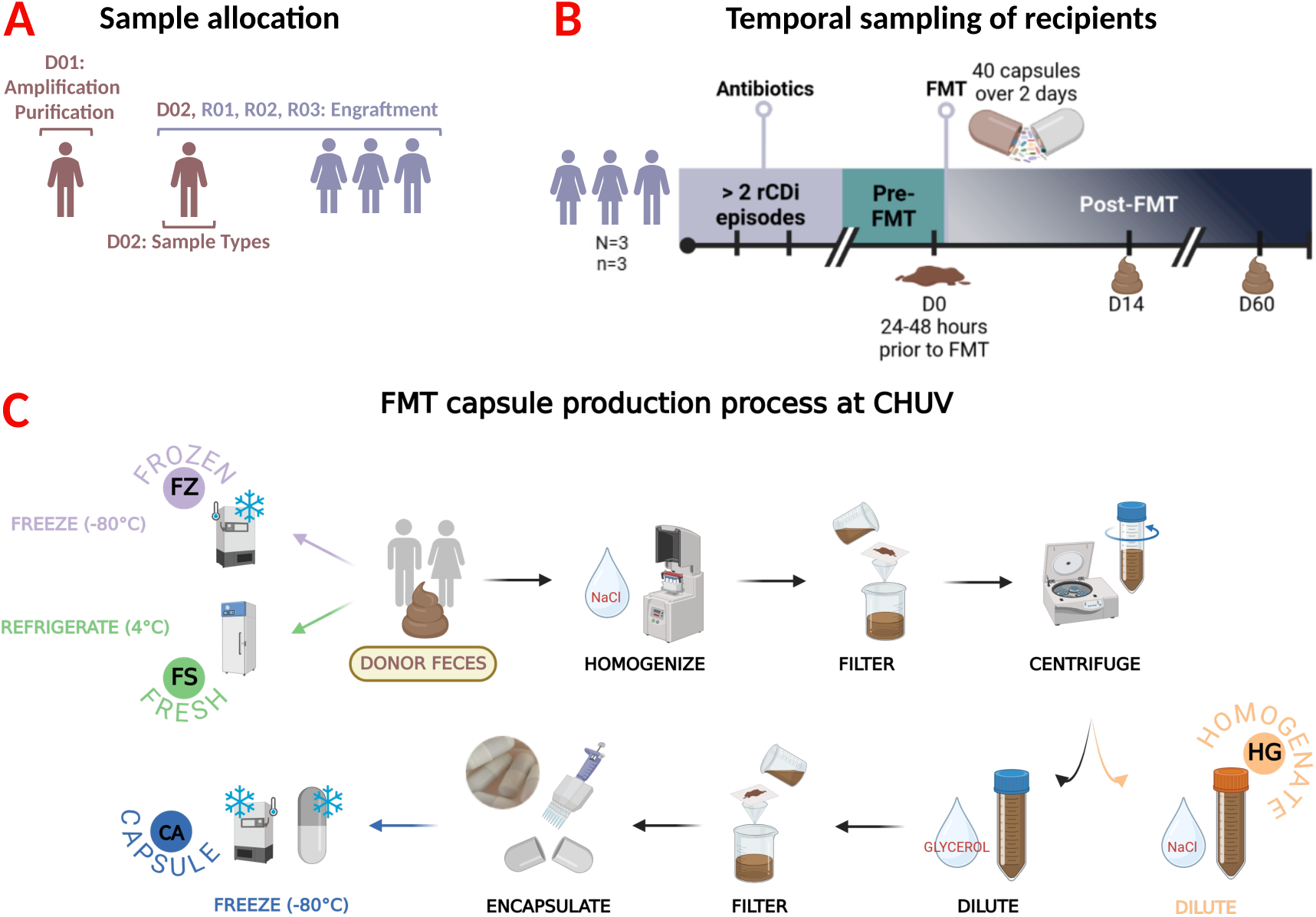
Recruitment of study participants and production of oral FMT capsules at CHUV. Donors who successfully passed a stringent screening following international guidelines were recruited. **(A)** Two FMT donors (D01 and D02) and three recipients were enrolled. Samples from D01 were used to evaluate the need for amplification and to compare spin-column versus magnetic bead purification. Samples from D02 were used to compare phage profiles across different sample types. D02 samples and the three recipient samples were further analyzed to assess donor phage engraftment. **(B)** Three patients experiencing rCDI were recruited. Samples were collected at baseline (pre-FMT), and at 14 and 60 days post-FMT to evaluate phage engraftment and temporal stability. **(C)** For capsule production, donor fecal samples were collected, then diluted, filtered, and centrifuged to remove large stool particles. The concentrated material was diluted in 0.9% NaCl to produce the homogenate, or suspended in glycerol (80 % m/v), followed by an additional filtration before encapsulation in size “00” capsules and storage at −80*^◦^*C. No cryoprotectant was used in fresh and frozen samples. Instead, fresh samples were stored at 4*^◦^*C, to provide a baseline reference, and frozen samples were stored at −80*^◦^*C. Figure created on biorender.com.

**Table 1:** Clinical characteristics of the donors and patients.

|  | Donors |  | Recipients |  |  |
| --- | --- | --- | --- | --- | --- |
|  | DO1 | DO2 | RO1 | RO2 | RO3 |
| **Characteristics** |  |  |  |  |  |
| Ethnicity | Caucasian | Caucasian | Caucasian | Caucasian | Caucasian |
| Diet | Flexitarian | Omnivore | Omnivore | Omnivore | Omnivore |
| Age (years) | 43 | 34 | 68 | 37 | 55 |
| Sex | Male | Male | Female | Female | Male |
| BMI | 20 | 24.8 | 30.8 | 23.8 | 27.1 |
| **CDI related** |  |  |  |  |  |
| Antibiotic treatment within 3 months prior to FMT (days) | - | - | Amoxicillin (7d) | Azithromycin (5d) | Cefepime (10d) |
| Number of CDI episodes | - | - | 3 | 3 | 2 |
| CDI treatment for the last episode <sup>1</sup> | - | - | Extended-pulsed fidaxomicin regimen | Vancomycin | Extended-pulsed fidaxomicin regimen |
| Duration of vancomycin FMT pre-treatment (days) | - | - | 12d | 11d | 12d |
| Clinical presentation of CDI <sup>2</sup> | - | - | Severe | Mild | Mild |
| Immunosuppression status | - | - | - | Yes | Yes |
| Notable comorbid condition <sup>2</sup> | - | - | Chronic obstructive pulmonary disease | Ileocecal Crohn's disease (on Vedolizumab) | Extranodal NK/T-cell lymphoma (on chemotherapy) |
| Clinical outcome <sup>3</sup> | - | - | Cured | Cured | Cured |

**Notes:**
All patients had undergone multiple rounds of antibiotics for previous episodes and were pre-treated with vancomycin (125 mg twice daily) while awaiting FMT to prevent further recurrences, which was discontinued 48 hours before the procedure. The FMT treatment regimen consisted of ingesting 2x20 FMT capsules within two days.
<sup>1</sup>Fidaxomicin extended-pulse regimen: oral film-coated tablets (200 mg twice daily for) the first five days, followed by one tablet every other day from day seven to day 25 <sup>21</sup>. Vancomycin: 125 mg four times daily for 10 days.
<sup>2</sup>The criteria for CDI severity and complications are defined in the ESCMID <sup>15</sup> or WHO <sup>22</sup> guidelines.
<sup>3</sup>Patients were considered cured if they remained symptom free 8 weeks post-FMT following ESCMID guidelines <sup>15</sup>.

### 3.2 Production process of oral FMT capsules

The process for producing frozen oral capsules was adapted from Youngster et al.^16^, with the only modification being an increase in glycerol content, and is illustrated in **Fig 1C**. To determine whether the pre-processing steps involved in producing the capsules affect the phage community, four sample types were evaluated from various stages of the production process: fresh (FS), frozen (FZ), homogenate (HG), and capsule (CA). Fresh samples were stored at 4*^◦^*C without any preservative media and provided a baseline phage profile. Frozen samples were stored at −80*^◦^*C prior to processing, to assess the impact of freezing on phages in the absence of cryoprotectants. Capsule production occurred within six hours of fecal collection; the material was diluted in 0.9% sodium chloride (NaCl), then filtered (diameter 125mm folded paper filter) to remove large undigested particles and centrifuged. The filtered material was either resuspended in NaCl to create the homogenate, or in glycerol (80% m/v) for further processing. The glycerol suspension underwent an additional filtration step prior to double encapsulation, with size “0” capsules enclosed within size “00” capsules, then frozen at −80*^◦^*C.

### 3.3 Viral particle enrichment and sequencing

The protocol for viral nucleic acid extraction and viral particles enrichment from human fecal samples was adapted from Monaco and Kwon^17^, with minor adjustments to increase DNA yield and improve sequence assembly quality. Briefly, the optimized protocol involves pulverizing thawed fecal material in cold saline magnesium buffer before low-speed centrifugation and filtration to remove large debris through 0.45 *µ*m (once) and 0.22 *µ*m (twice) polyethersulfone membrane filters. To enrich and purify viral particles, samples were treated with lysozyme, chloroform, DNase, SDS/CTAB, proteinase K, and phenol:chloroform:isoamyl alcohol.

To investigate whether DNA amplification was necessary, unamplified and amplified (GenomiPhi V2 DNA Amplification Kit, Cytiva) samples from D01 were compared for DNA yield and quality of genome assembly following sequencing. GenomiPhi employs multiple displacement amplification (MDA) to amplify DNA under isothermal conditions by random priming using the highly replicative *ϕ*29 polymerase. For this initial round of sequencing, aimed to evaluate and improve the protocol, DNA was purified using the DNeasy Blood and Tissue Kit (Qiagen, Germany). Library preparation was performed with Nextera XT, and sequencing was conducted on an Illumina MiSeq platform. Twelve shotgun libraries produced 35,681,410 quality-filtered trimmed reads, with a median read count of 3,054,742 per sample.

In subsequent sequencing rounds, DNA was purified using AMPure XP (Beckman Coulter, USA). DNA Prep kit (Illumina, USA) was used for library preparation as it is better suited for low DNA concentrations. These 25 libraries were sequenced on an Illumina NextSeq with a P1 cassette to generate a higher number of reads, resulting in 408,742,766 quality-filtered trimmed reads, with a median read count of 16,194,692 per sample.

### 3.4 Sequence analysis

Taxonomic classification of reads was performed using Kraken2 (v2.1.3) (Retrieved June 5, 2024, standard database, IndexZone. Quality-controlled and trimmed reads (BBMap v39.06 using the following settings: ktrim=r tpe=f tbo=f qtrim=rl trimq=15) were assembled into contigs either individually or across all samples (referred to as the co-assembly) using MEGAHIT (v1.2.9; default parameters). Assembly metrics and coverage were evaluated using QUAST (v5.2.0) and CoverM (v0.7.0), respectively. The completeness of viral genomes was assessed with CheckV (v0.7.0, database v1.5) and VirSorter2 (v2.2.4). High-quality viral genome (n=332) were extracted from the co-assembly. Sample reads were mapped to these genomes, and abundances were normalized for genome length and sequencing depth to generate sequence counts per genome size per million (CPGM) values. Predictions of phage lifecycle, taxonomic classification, and bacterial hosts were performed using the Unified Human Gut Virome Catalog and toolkit (UHGV, v0.0.1).

### 3.5 16S rRNA metagenomic processing

A pea-sized quantity of stool suspended in 1 mL phosphate-buffered saline (PBS) was centrifuged (10 min x 500 rcf) to pellet large stool particles. The supernatant was collected and concentrated (10 min x 13,000 rcf). MagNA Pure (Roche, Switzerland) was used to extract the DNA, and the V3-V4 region of the 16S rRNA was sequenced using MiSeq platform (Illumina, USA). The reads were processed using zAMP (v0.9.19)^18^, an amplicon-based metagenomic pipeline consisting of paired-end assembly, removal of adapters and primer sequences (Cutadapt), length trimming, amplicon sequence variant (ASV) denoising (DADA2), and taxonomic assignment (EzBioCloud, 2018.05 release, pre-processed^19^). Reads were rarefied to approximately 50,000 per sample, with singletons and doubletons removed. Correlations between bacterial and phage profiles were assessed by comparing the relative abundance of bacterial hosts, as determined by 16S rRNA sequencing and host predictions from UHGV.

### 3.6 Statistical analysis and visualization

Statistical analyses were conducted on RStudio (v4.2.1) using packages including vegan, ggpubr MicrobiotaProcess, pairwiseAdonis, and lme4. Significance between DNA yield and raw read counts was assessed using a t-test. Visualization of the viral co-assembly was performed using Anvi’o (v8)^20^, with additional visualizations generated using ggplot2 and ComplexUpset. For all preservation and engraftment analyses, genomes with a breadth of coverage *<* 80% were considered absent. To evaluate the effect of sample types on alpha diversity metrics (Chao1, Pielou, and Shannon indices), a Tukey’s post-hoc test was applied to a linear mixed-effects model (no missing data). Bray-Curtis dissimilarity was used to assess differences in viral compositions between groups. For analyses comparing sample types, donations were treated as a blocking variable. Similarly, when comparing recipients, timepoints within each recipient were treated as a blocking variable to account for variation. Given the likely correlation between phage and bacterial profiles^10^, the depth of phages associated with each predicted bacterial host was used as a proxy to estimate the bacterial abundance.

## 4 Results

### 4.1 Characteristics of the study population

The characteristics of the two FMT donors and three recipients recruited are detailed in **Table 1**. Donors were, on average, younger (38.5 years) and had a lower body mass index (BMI, 22.4) compared to patients (53.3 years, BMI 27.3), Both donors were male, one following a flexitarian diet and the other an omnivorous diet. Two of the three patients were female, and all adhered to an omnivorous diet. Recipients had received multiple courses of antibiotics for prior CDI episodes and were pre-treated with vancomycin, which was stopped 48 hours before FMT. Both R01 and R02 experienced three rCDI episodes. R01 suffered severe episodes, characterized by fever, a white blood cell count *≥*15,000 cells/*µ*L, and underlying chronic obstructive pulmonary disease. R02 was additionally treated with ENTYVIO for Crohn’s disease. R03 developed two rCDI episodes and was immunosuppressed due to extranodal NK/T-cell lymphoma undergoing chemotherapy.

### 4.2 Protocol optimization for viral particle sequencing

The DNA yield from viral particles was low due to their small genome size and limited sample availability. In an attempt to enhance DNA concentration for sequencing, we employed random primer amplification using *ϕ*29 polymerase. To evaluate the impact of DNA amplification on both the quantity and quality of sequencing reads and assembled contigs, we compared six paired D01 samples prepared without amplification to those amplified using the GenomiPhi V2 DNA Amplification Kit (Fig 2). As expected, amplified samples resulted in a significantly higher yield of DNA (ca. 2400*×*, *p <* .0001) and raw reads (*p <* .0001) (**Fig 2A-B**). The majority of reads in both amplified and unamplified samples could not be classified by Kraken2, with only a small fraction being classified as viral or bacterial. On average, unamplified samples had more reads classified as viral (ca. 5*×*, *p* = 0.0006) (data not shown). Interestingly, while amplified samples generated more raw reads, unamplified samples exhibited a larger assembly length and a greater number of contigs *>* 1 kilobases (kb) (**Fig 2C**). In contrast, the majority of contigs assembled from amplified samples were *<* 1 kb. This trend was also reflected in the quality assessment from CheckV and VirSorter2. Unamplified samples consisted of a higher proportion of contigs categorized as medium, high, or complete quality, as well as a greater number of complete genomes (**Fig 2D**, data for VirSorter2 not shown). Based on these results, subsequent experiments were conducted without amplification to generate a greater number of higher-quality contigs and more complete genomes.

**Figure 2:**
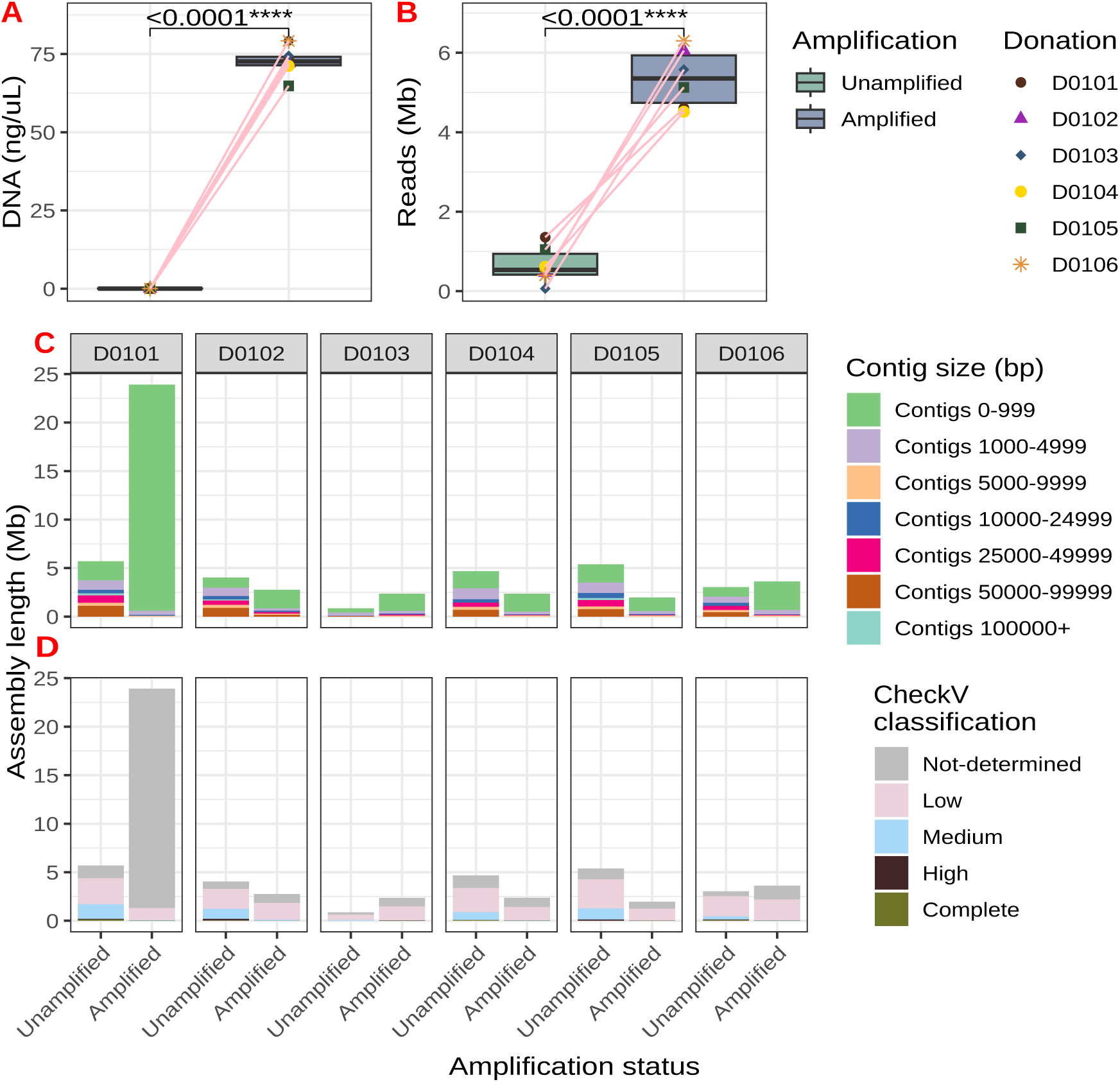
Assessing the impact of amplification on the metagenomic analysis of the human fecal phageome. The quality of assembled genomes was evaluated to determine whether amplification of viral nucleic acid was necessary. **(A)** Unamplified samples exhibited significantly lower DNA concentration prior to library preparation, **(B)** which resulted in fewer reads. **(C)** However, unamplified samples yielded more contigs of larger sizes, while the majority of amplified contigs were *<* 1 kb. (**D**) Additionally, unamplified samples produced a greater number of higher-quality contigs. Based on these findings, subsequent experiments proceeded without amplification.

### 4.3 Selection of high-quality viral contigs from co-assembly

Kraken2 was used to classify sequences obtained from donor samples and identify the proportion of viral reads. A median proportion of 35.6% *±* 11% and 17.7% *±* 33.1% of the quality-filtered and trimmed reads were identified as viral in the donor and recipient samples, respectively. Most reads were unclassified (56.2% *±* 10.6% and 69% *±* 28.9%, respectively), which could potentially include unidentified phages. Only a small proportion of reads were identified as human or bacterial.

To obtain a reference phageome across all 25 donor and recipient samples, sequencing data from 16 samples collected at various production stages from five donations of one healthy donor, together with nine samples from three FMT recipients (before, and 14 and 60 dp-FMT) were co-assembled to increase coverage depth and breadth, as well as the number of genomes recovered. A total of 157,137 contigs were obtained, of which 84% were *<* 1 kb and 0.79% *>* 10 kb. The total cumulative assembly length was 151.8 million bp (Mb).

Only 333 contigs (0.21%) were classified to be complete, high-quality, or medium-quality by CheckV, and full or partial genome by VirSorter2 (**Fig 3A**). To focus on high-quality viral contigs, one lower-quality contig that had a coverage *<* 80% was excluded. The remaining 332 viral contigs ranged in size from 2,561 to 344,393 bp, collectively representing 10.5% of the total co-assembly (15.9 Mb) (**Fig 3B-C**). Based on the UHGV database, contigs were assigned to 18 classifications, mostly at the class level. Of these contigs, 83.4% were classified as belonging to Caudoviricetes, while *Microviridae* (6.3%) and *Salasmaviridae* (3.6%) were the most abundant families. Most phages were predicted to have a temperate (65%) or virulent (31%) life cycle. Predicted bacterial hosts included Clostridia (9%), Clostridiales (5.7%), *Lachnospiraceae* (8.1%), and *Oscillospiraceae/Ruminococcaceae* (5.7%).

**Figure 3:**
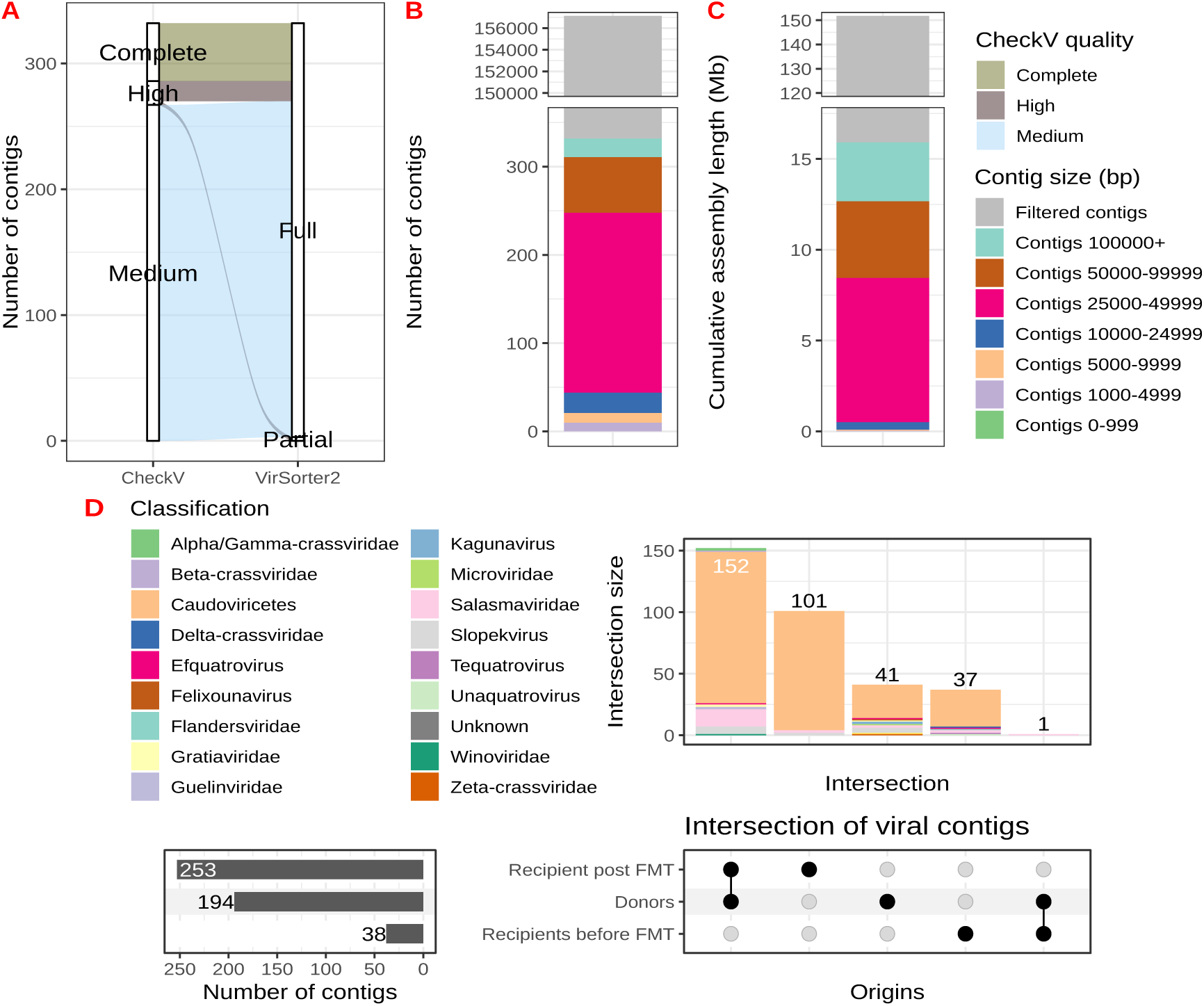
Quality assessment of the viral co-assembly. All samples (N=25) were co-assembled to obtain a reference phageome. **(A)** High-quality viral contigs (n=332), classified as complete, high, or medium quality by CheckV, and full or partial genome completeness by VirSorter2, were extracted. Colors represent the quality classification by CheckV. **(B-C)** The high-quality viral contigs range from 2,561 to 344,393 bp, with their cumulative assembly length accounting for 10.5% of the total co-assembly. The colors indicate the distribution of contig sizes. **(D)** Of the 332 phages, the highest prevalence was observed in recipients after treatment, followed by donors and recipients before FMT. The phages are classified into 18 taxa by UHGV, with Caudoviricetes being the most abundant phage. Colors represent the predicted phage classifications.

### 4.4 Preservation of phages in capsules

To assess whether the capsule production process, which includes filtration, centrifugation, and storage in preservative media, alters the phage profile, sixteen samples from five donations were collected at various stages of the production process: fresh, frozen, homogenate, and capsule (**Fig 4A-C**). Fresh samples provided a baseline phage profile. Viral nucleic acid was successfully extracted from all sample types in comparable quantities. Hierarchical clustering of donor samples based on the presence and absence of the 332 high-quality viral genome sequences revealed that capsules cluster more closely with their fresh counterparts than frozen and homogenate samples **(Supp Fig 1**). When considering abundances rather than presence and absence, samples from the same donation clustered more closely, with capsules again positioned closest to their fresh counterparts, suggesting the proportion of phages remained consistent across different processing methods. The donor signature was highly conserved, as evidenced by the consistent presence and abundance of contigs across donations and sample types. To assess the changes in phage composition within sample types, alpha diversity metrics such as Chao1 (richness), Pielou (evenness), and Shannon (richness and evenness) indices were used (**Fig 4B**). A Tukey’s post-hoc test applied to a linear mixed effects model revealed a significant reduction in Chao1 diversity in the frozen (*p* = .03) and homogenate (*p* = .03) samples compared to the fresh, and a reduction in Shannon index in homogenate compared to fresh (*p* = .04). Phage composition between sample types was compared using Bray-Curtis dissimilarity (**Fig 4C**). A multivariate homogeneity of group dispersions (*p* = .72) and pairwise permutational multivariate analysis of variance (PERMANOVA) using distance matrices (*p >* .05) did not identify any significant difference between sample types.

**Figure 4:**
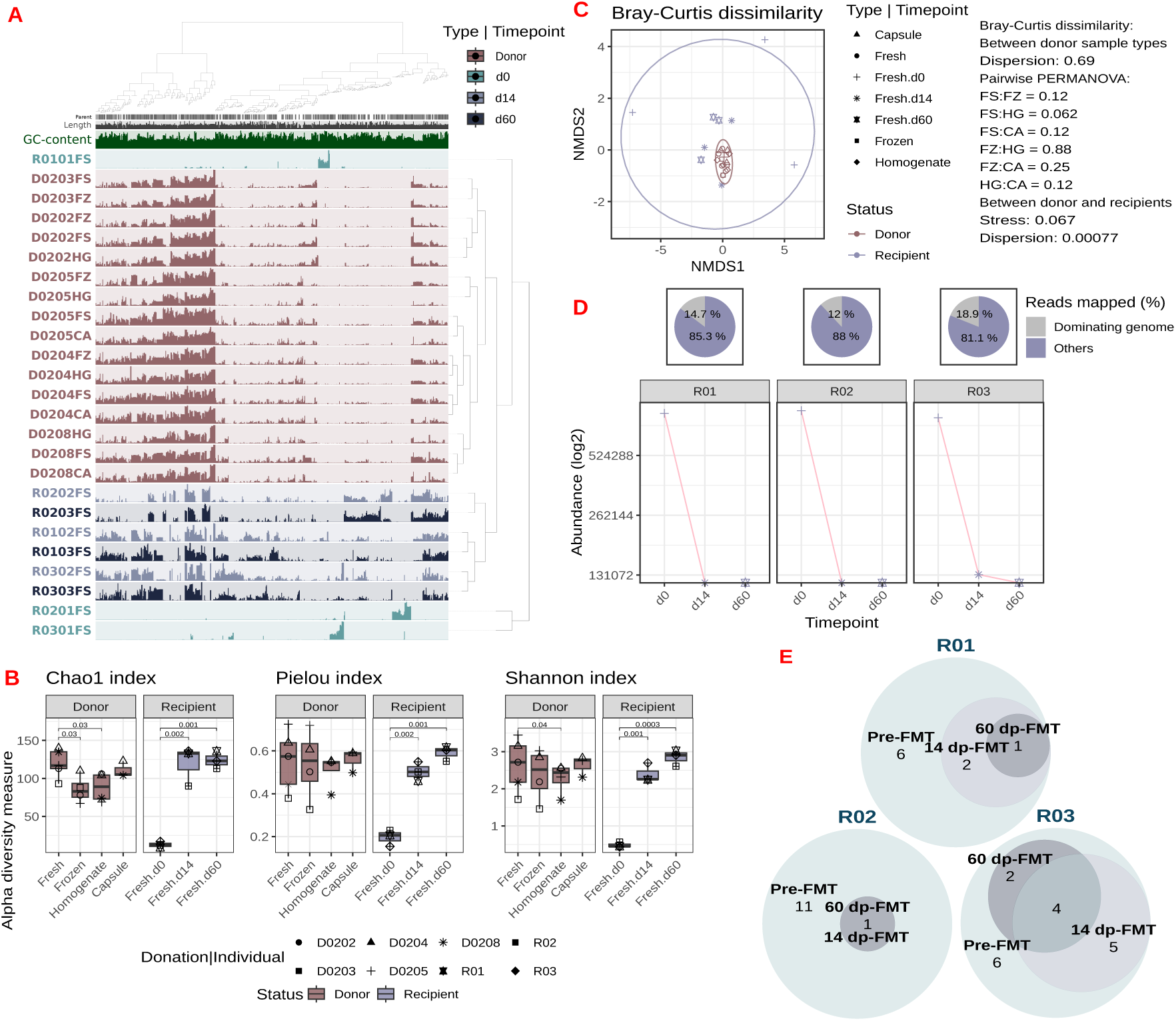
Preservation of phages across sample types and engraftment of donor-derived phages. Phage profile based on the abundance of 332 high-quality genomes. **(A)** Donor samples clustered within their respective donation, irrespective of sample type, and capsules clustered most closely with their fresh counterparts. Pre-FMT samples were the most distinct, while post-FMT samples clustered closely together, with the 60-day profiles appearing stable. Engraftment of donor-derived phages was observed. Sample labels indicate the type: FS (baseline, fresh), CA (capsule), FZ (frozen), and HG (homogenate). The colors represent the donors (red), recipients pre-FMT (light blue), 14 (medium blue), and 60 dp-FMT (dark blue). Phages were considered present if the breadth of coverage was *≥* 80%. Phage genomes were split into 20,000 bp fragments and clustered based on coverage, with samples ordered according to sequence composition and differential coverage using Euclidean distance and Ward’s method for hierarchical clustering. Within each sample (row), the mean coverage of each genome is displayed (generated using Anvi’o^20^). **(B)** Chao1 (richness), Pielou (evenness), and Shannon (richness and evenness) indices were assessed to evaluate the change in composition across samples. Capsules showed comparable alpha diversity to fresh samples, as indicated by a linear mixed-effects model. Recipients before FMT exhibited a significantly lower alpha diversity compared to their post-FMT profiles. The colors indicate the donor or recipient status, while shapes represent the donation or the individual the sample was obtained. **(C)** Bray-Curtis dissimilarity was used to compare the composition between groups. Donor samples cluster closely together, with no significant compositional differences between sample types. In contrast, recipients before FMT were highly distanced, while post-FMT samples closely resembled the donor profile. Colors indicate donor or recipient status, while shapes correspond to the sample type or timepoint. **(D)** The viral profiles of recipients prior to FMT exhibit low viral diversity, marked by the dominance of a single phage per individual. The line plot displays the temporal changes in the normalized abundance (CPGM) of dominant phage for each recipient, while the pie chart shows the proportion of reads mapped to the dominant phage (purple). **(E)** Several phages across the three recipients (11.4%) were identified to be intrinsic to the recipients; however, most were lost following FMT. The stability of these intrinsic phages after FMT is illustrated. Colors indicate the number of intrinsic phages present pre-FMT (lightest), 14 (medium), and 60 dp-FMT (darkest).

### 4.5 Transfer of donor-derived phages to recipients via FMT capsules

The profiles of donors and recipients were evaluated to confirm the engraftment of donor phages (**Fig 4A-C**). Recipients before FMT exhibited a distinctly separate profile from all other samples, marked by low phage diversity (**Fig 4A**). On the contrary, profiles after FMT, clustered by the recipient, and were characterized by increased diversity, with a stable composition observed at 60 dp-FMT. Moreover, the emergence of new phages was observed. All alpha diversity metrics in recipients prior to FMT were significantly lower compared to 14 (*p <* .002) and 60 (*p <* .001) dp-FMT (**Fig 4B**). However, no differences between post-FMT profiles were observed (*p >* .05). These findings were also visually reflected by the Bray-Curtis NMDS plot, where the pre-FMT samples were clearly separated from donor and post-FMT profiles, while post-FMT samples clustered more closely to donor samples, suggesting the engraftment of donor-derived phages (**Fig 4C**). An analysis of multivariate homogeneity of group dispersions between sample type and timepoint was significant (*p* = .0057); therefore, a permutational multivariate analysis of variance using distance matrices was not performed. However, a stress value of 0.088 suggests NMDS provides a reliable representation of the data.

Of the 332 phages, 46% were present in donors and recipients after treatment (**Fig 3D**). None of the identified phages were shared between donors and recipients pre-FMT, while 101/332 (30%) were identified only in recipients after treatment. All profiles were dominated by Caudoviricetes class, with donors exhibiting the most diverse phage community (14/18 classifications, 77.8%), followed by post-FMT recipients (9/18, 50%). Pre-FMT recipients displayed the least diversity (7/18, 38.9%).

The relationship between the depth and breadth of coverage for each contig was explored. This revealed that each recipient before FMT had a distinct dominant phage, which represented the majority of their respective reads mapped to the 332 high-quality viral genomes (81-88%) (**Fig 4D**). Although these phages could not be classified at the species level, they were predicted to be virulent phages, their large sizes suggesting they are likely complete genomes. In R01, a Caudoviricetes (58,992 bp) predicted to infect Proteobacteria was predominant, with 85.3% of all viral reads mapping to this phage. In R02, the most abundant phage was a *Felixounavirus* (89,440 bp) predicted to infect *Escherichia*, with 88% of reads mapped to it. Finally, R03 was dominated by a *Delta-crassviridae* (99,648 bp) predicted to infect Bacteroidota, which represented 81.1% of all mapped reads. After FMT, these phages were undetectable in two recipients and decreased by 13% in the third recipient at 14 dp-FMT, undetectable by day 60.

The temporal changes in recipient-specific phages were also evaluated (**Fig 4E**). Across the three recipients, 38 of the 332 phages (11.4%), with a minimum coverage threshold of 80%, were detected prior to FMT (**Fig 4D**). Most of these 38 phages (95%) were unique to each recipient and were subsequently not detected after FMT, a trend consistently observed across all three individuals. In R01, three of the nine contigs persisted at day 14, but only one remained stable at day 60. In R02, only one of the 12 pre-FMT contigs remained stable through 14 and 60 days after FMT. Four of the 17 contigs were stable at both 14 and 60 dp-FMT in R03, while five were not observed after day 14, and two re-emerged after 14 days. Overall, most of the phages intrinsic to recipients were no longer detectable following FMT.

### 4.6 Correlation between predicted phage host and observed bacterial abundance

To explore the correlation between the phage and bacterial communities, 16S rRNA amplicon sequencing was additionally performed in the initial stool sample. Using host classifications predicted from the UHGV catalog, the relative abundances of phages were compared to their corresponding bacterial hosts identified according to 16S rRNA sequencing. An inverse correlation between the phage and bacterial abundances was observed in pre-FMT samples (Fig 5). Following FMT, samples exhibited increasing complexity and diversity in both the phage and bacterial profiles, indicating a transition to an eubiotic-like state.

**Figure 5:**
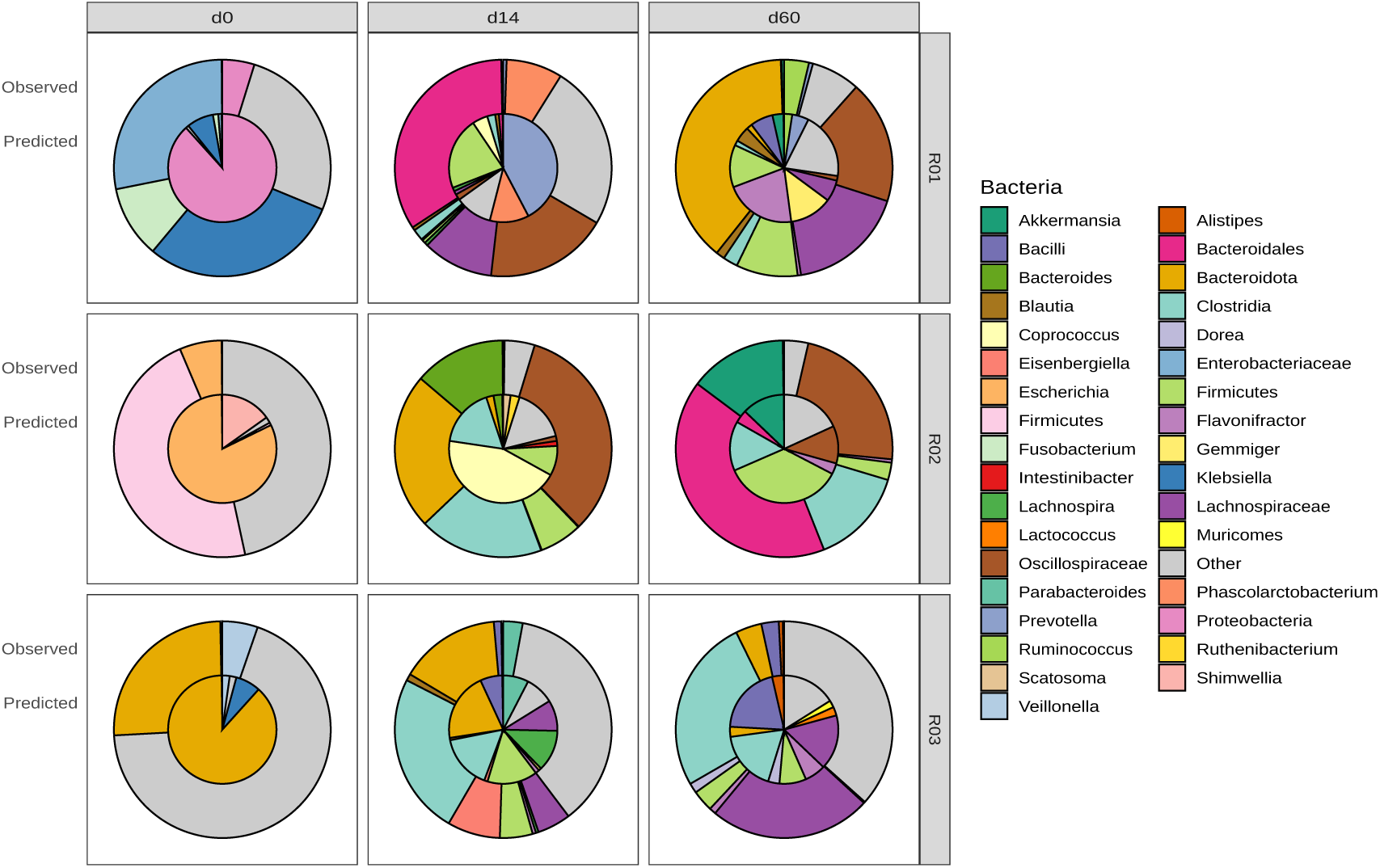
Correlation between phage and bacterial profiles. The relative abundance of the most dominant bacterial taxa associated with phages predicted by the UHGV catalog (inner ring) was compared to the corresponding bacterial taxa identified by 16S rRNA amplicon sequencing (outer ring). Colors represent bacterial taxa across various taxonomic ranks.

## 5 Discussion

Recent studies demonstrated that FVT effectively restores normal stool habits and eliminates symptoms of CDI, highlighting a potential role for phages and/or bacterial metabolites^11,14^. However, these studies focused on the phage community extracted directly from fresh stool material. At the Lausanne University Hospital, rCDI patients are treated with frozen oral FMT capsules, which undergo various pre-processing steps that may alter the phage profile. To investigate whether the phage component of these capsules could contribute to FMT success, the phage profiles of individuals treated with capsules were characterized to assess the impact of pre-processing steps during capsule production, and to analyze the engraftment of donor-derived phages as well as their temporal stability.

During the protocol development, we observed that MDA disproportionately amplified small contigs (*<* 1 kb), resulting in a higher proportion of low-quality classifications and incomplete genomes compared to unamplified samples, as classified with CheckV and Virsorter2. These results contrast the frequent use of MDA in protocols for metagenomic analysis of the human fecal phageome^17,23,24^, Appendix C calling for a change in practice. While MDA is effective for amplifying ultra-low amounts of DNA, it introduces several biases including chimera formation, preferential amplification of small single circular ssDNA genomes (due to the absence of a denaturing step), biases against DNA with low and high GC content, and uneven amplification of linear dsDNA genomes, despite pooling independent reactions^23,25,26^, Appendix C. Furthermore, evidence that MDA introduces biases that can alter the phage profile has been emerging. For example, decreased phage diversity and reproducibility^27^, while over- and under-representation of the OTUs relative to the controls was reported^25^. Furthermore, Reyes et al.^24^ demonstrated that although a greater proportion of sequences from unamplified datasets were kept in MDA-amplified datasets, fewer sequences from the MDA-amplified datasets were recovered in the unamplified ones, highlighting the biases introduced by amplification. Given these findings, and in agreement with other studies^24^, Appendix C we omitted the MDA step from our final protocol to minimize bias and improve the quality of viral metagenomic data.

From the co-assembly, 332 high-quality phage genomes (*≥* 80% coverage) were identified, with the majority classified as Caudoviricetes by UHGV. This finding aligns with previous studies suggesting that *Caudoviridae*, now reclassified as Caudoviricetes, and *Microviridae* are the most abundant members in the gut phageome^7^. The majority of phages were predicted to have temperate life cycles. Unlike lytic phages, which immediately lyse their host, temperate (lysogenic) phages are able to integrate into the bacterial genome as latent prophages, replicating alongside the bacterial host. Under stressful conditions, they can enter a lytic cycle, leading to host cell lysis^28^. Temperate phages are believed to play a role in microbiota stability, through mechanisms such as horizontal gene transfer and the regulation of bacterial populations^28,29^, however, their precise role remains to be understood.

To assess whether the capsule production altered the phage profile, samples were collected at different points throughout the production process. The presence of viral contigs in capsules confirmed that phages were preserved. It is well established that individuals possess their own stable and unique microbial profiles^30,31^, Appendix C. Accordingly, samples clustered according to their respective donations, with capsules closest to their fresh counterparts. This indicates that the pre-processing steps did not significantly alter the phage community, and donation-specific variations were retained. However, when examining the presence and absence of phages, frozen and homogenate samples clustered together. Additionally, a post-hoc Tukey’s test evaluating the alpha diversity, identified a significant reduction in Chao1 index in the frozen or homogenate samples, and a decrease of Shannon index in the frozen, compared to the fresh samples. The homogenate differed from the capsule by being suspended in NaCl instead of glycerol, while the frozen samples lacked any cryoprotectant. NaCl maintains osmotic balance and cell integrity; however, unlike glycerol, it is not a preservative. Glycerol prevents ice crystal formation by bonding with water molecules^32^, Appendix C, suggesting its suitability for preserving both bacteria and phages. Additionally, beta-diversity analysis confirmed that the overall composition remained comparable across sample types, suggesting that the pre-processing steps did not significantly alter the phage profile.

To evaluate whether phages successfully engrafted in recipients, samples prior to and post-FMT were collected. Hierarchical clustering revealed that the pre-FMT profiles were distinctly different from post-FMT profiles. Post-FMT profiles clustered more closely with the donor profile. Moreover, alpha diversity indices within donor or post-FMT profiles were comparable. Finally, beta diversity analysis revealed that post-FMT and donor samples clustered together, while pre-FMT profiles remained distant. Together, these findings indicate the successful engraftment of donor-derived phages in recipients.

Prior to treatment, the viral community in each recipient was dominated by a distinct single contig, which accounted for 81-88% of their total reads. This, along with the low Chao1 index, indicates that the pre-FMT profile is characterized by low viral diversity, a finding consistent with previous studies^11,33^. This low diversity may be attributed to the CDI and subsequent vancomycin treatment prior to FMT Appendix C, which in turn could have promoted more effective engraftment by reducing niche and nutrient competition.

As bacterial and phage profiles are linked, it is important to consider how changes in one composition alter the other; the loss of bacterial diversity may influence the viral composition as well^10^, Appendix C. Post-FMT profiles at days 14 and 60 were compared to evaluate the stability of the phage community after treatment. Before FMT, recipients exhibited low *Microviridae* abundance; however, after FMT, the transfer of phages from donors resulted in an increase in *Microviridae*, a pattern consistent with the findings from Zuo et al.^11^. Profiles at 14 and 60 dp-FMT clustered by the recipient, indicating the phage community remained stable over time in a given subject, while maintaining inter-individual variation. This intra-individual stability was further supported by the comparable alpha and beta diversity profiles between the two timepoints. Post-FMT profiles also reveal the emergence of new phages not detected in donors and recipients before FMT. These may have been present initially at low abundance in the recipient or the donor, but classified as absent due to the coverage threshold *<* 80%, or they could have been introduced through sources other than FMT, such as dietary intake. Phages have been isolated from fermented food, such as kimchi, cheese, and sauerkraut, and other products, such as toothpaste, stevia, and honey may also induce prophage activation Appendix C.

Comparison of the phage and bacterial profiles reveals an inverse correlation in pre-FMT profiles, an observation also supported by others^34,35^, and supports the “killing the winner” hypothesis, that phages may be playing a role in promoting community balance^36,37^. An increase in phage and bacterial diversity post-FMT was also observed, highlighting the transition to a healthier state^38–40^.

While these insights into the phageome following FMT in rCDI patients align with previous research, we acknowledge that the sample size may affect the statistical power and generalizability of the results. Finally, most of the contigs (99.79%) were excluded to focus on high-quality contigs, which may also impact our interpretation of the true phage diversity in the gut. However, this filtering was needed to avoid spurious hits, and our results capture the most abundant and diverse phages in these inherently complex communities.

## 6 Conclusion

In this study, we investigated whether the production process of CHUV’s oral FMT capsules preserves the phage composition of donor material and whether these phages successfully engraft in recipients. Our findings reveal that the encapsulation process maintains the integrity of the phage community and that phage engraftment occurs post-transplantation, as reflected by a notable increase in phage diversity. Many of these engrafted phages are predicted to infect key gut commensals, suggesting that they may play an active role in reshaping the host microbiome. These results point to a previously underappreciated contribution of the virome to FMT outcomes. To fully unravel the role of phages in therapeutic success, future studies should incorporate larger and more diverse patient cohorts, include individuals with FMT failure, and conduct comparative trials between FMT and FVT. Longitudinal monitoring integrating phage, bacterial, and metabolomic profiling will also be essential to decode the complex dynamics underlying microbiome-based interventions.

## Acknowledgments

The authors would like to thank Rizlène Dira, Olga Martin, the CHUV team for Fecal Microbiota Transplantation, the sequencing platform of the DMLP, Nathalie Bonvin, and Sabrina Pereira Pipa for their assistance in recruiting participants, collecting information, and preparing the samples.

## 8 Data availability

All data generated in this study have been deposited in the European Nucleotide Archive (ENA) at EMBL-EBI under accession number PRJEB97162. The R scripts used for statistical analyses and figure generation are publicly available on Zenodo at https://doi.org/10.5281/zenodo.17290109.

## 9 Declaration of generative AI and AI-assisted technologies in the writing process

During the preparation of this work the authors used OpenAI’s ChatGPT and Grammarly in order to assist with the grammatical refinement of the paper. After using this tool/service, the authors reviewed and edited the content as needed and takes full responsibility for the content of the publication.

## A Supplementary Methods

### A.1 Recruitment process

Briefly, eligibility criteria for FMT donors included: good general health, age 18–50 years, no significant personal or family history of medical conditions, no recent use of medications affecting the gut microbiota, regular intestinal transit, and stool consistency corresponding to 2–5 on the Bristol Stool Scale. Recipients with missing samples at any timepoint were excluded.

## B Supplementary Figures

**Supp Fig 1:**
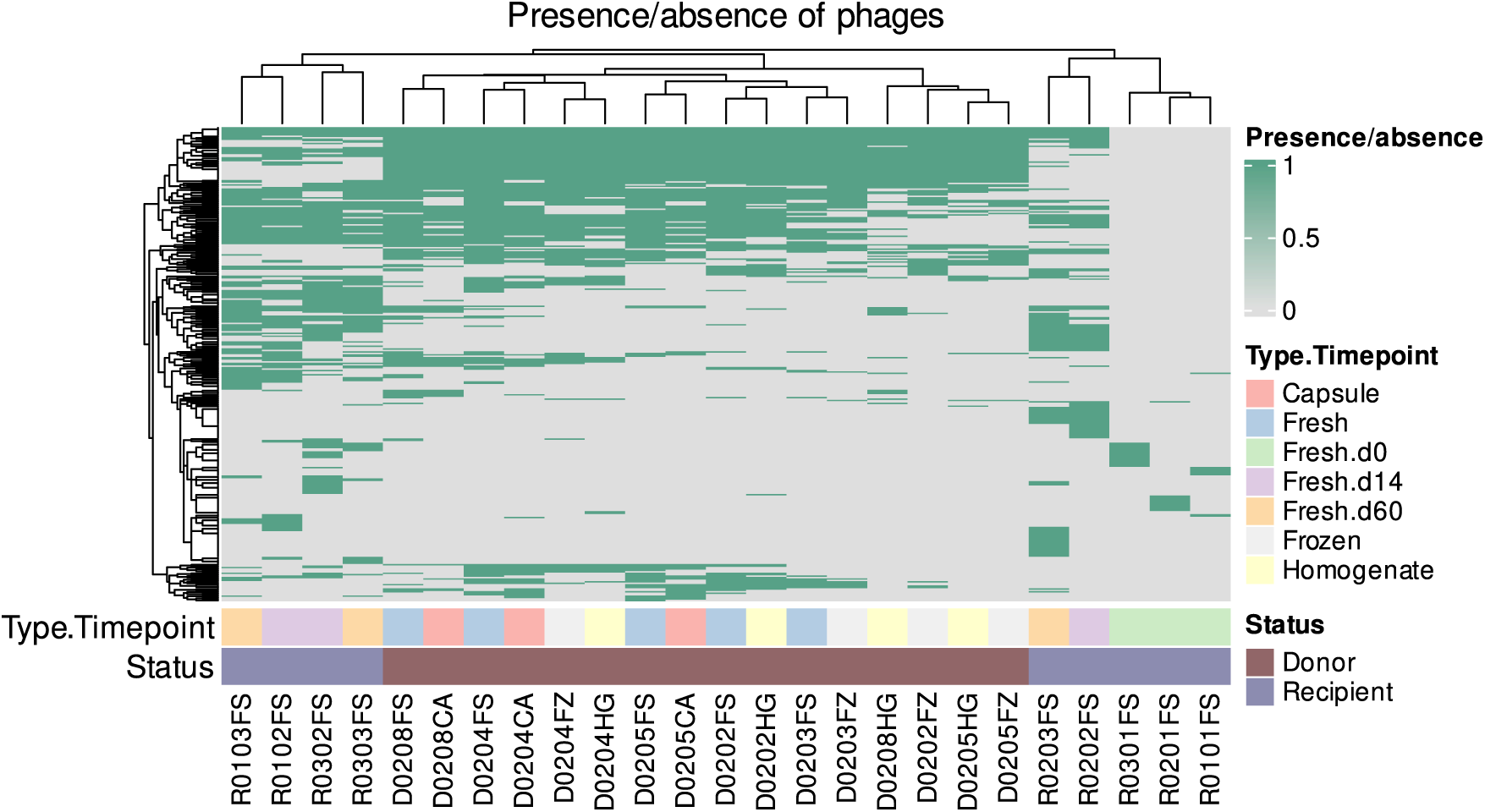
Presence and absence of phages. Donor samples cluster together, while pre-FMT samples cluster furthest away. Post-FMT samples group by recipient. Genomes with a sequence coverage *≥* 80% were considered present. Type.Timepoint indicates the donor sample type or recipient timepoint, while status specifies whether the individual was a donor or recipient.

## C Supplementary References

Criteria for screening FMT donors according to international guidelines are detailed in several key publications^15^,^41^-^44^. Cryoprotectants can impact the composition and stability of phages^32^,^45^-^48^.

Multiple studies have employed MDA for human fecal phageome metagenomics^17^,^23^,^24^,^49^-^54^, although others have highlighted the limitations and biases introduced by MDA, including amplification bias, chimera formation, and GC content effects^23^,^25^,^26^,^55^,^56^. Studies conducted without MDA provide complementary perspectives on fecal phageome composition^24^,^57^-^59^.

Several studies have established that individuals possess stable and unique microbial profiles over time^30^,^31^,^60^-^62^. In the context of FMT, CDI and vancomycin treatment prior to transplantation reduce microbial diversity^63^-^65^, which may influence engraftment dynamics. Viral and bacterial profiles have been shown to correlate in the gut community^10^,^66^,^67^. Certain fermented foods, such as kimchi, cheese, and sauerkraut, and other products, such as toothpaste, stevia, and honey may also induce prophage activation^68^-^70^.

